# A lung targeted miR-29 Mimic as a Therapy for Pulmonary Fibrosis

**DOI:** 10.1101/2021.12.22.473724

**Authors:** Maurizio Chioccioli, Subhadeep Roy, Kevin Rigby, Rachel Newell, Oliver Dansereau, Linda Pestano, Brent Dickinson, Farida Ahangari, Gisli Jankins, Stewart Iain, Guari Saini, Simon R Johnson, Rebecca Braybrooke, Jose Herazo-Maya, Nachelle Aurelien, Guying Yu, Maor Sauler, Giuseppe DeIuliis, Rusty L Montgomery, Naftali Kaminski

**Affiliations:** Pulmonary, Critical Care and Sleep Medicine, Yale School of Medicine, New Haven, CT, USA; miRagen Therapeutics, Inc, Boulder, CO, USA; University of South Florida, Tampa, FL, USA; Hospital Medicine, Weill Cornell School of Medicine, NY, USA; Henan Normal University: Xinxiang, CN; National Heart and Lung Institute, Imperial College London, London, UK; Division of Respiratory Medicine and Respiratory Research Unit, University of Nottingham, Nottingham, UK

**Keywords:** Idiopathic Pulmonary Fibrosis, microRNA, miR-29, RNA therapies

## Abstract

microRNAs are non-coding RNAs that negatively regulate gene networks. Previously, we reported a systemically delivered miR-29 mimic MRG-201 that reduced fibrosis in animal models, but at doses prohibiting clinical translation. Here, we generated MRG-229, a next-gen miR-29 mimic with improved chemical stability, conjugated with the internalization moiety BiPPB (PDGFbetaR-specific bicyclic peptide). In TGF-b-treated human lung fibroblasts and precision cut lung slices, MRG-229 decreased COL1A1 and ACTA2 gene expression and reduced collagen production. In bleomycin-treated mice, intravenous or subcutaneous delivery of MRG-229 downregulated profibrotic gene programs at doses more than ten-fold lower than the original compound. In rats and non-human primates, and at clinically relevant doses, MRG-229 was well tolerated, with no adverse findings observed. In human peripheral blood decreased mir-29 concentrations were associated with increased mortality in two cohorts potentially identified as a target population for treatment. Collectively, our results provide support for the development of MRG-229 as a potential therapy in humans with IPF.

**One Sentence Summary:** One Sentence Summary: A stabilized, next-generation miR-29 mimic has been developed that demonstrates efficacy at commercially viable doses with a robust safety margin in non-human primates.

## Introduction

Most human diseases are not treatable with compounds that target a single protein or enzyme: hence, broad-acting therapies that can mitigate or reverse disease-associated molecular and cellular processes are urgently needed. In the search for modifiers of complex disease phenotypes, oligonucleotide (ON) technologies, with their rich array of modalities and capabilities for gene silencing, gene activation and splice modulation, are of particular interest ^1^. Progress towards their clinical translation into approved, effective and safe therapies continues to be made, as evidenced by FDA-approval of single strand steric block (exon-skipping) antisense ON drugs Exondys51 and Spinraza to treat patients with Duchenne’s muscular dystrophy and double strand siRNA drug Onpattro, which binds to and degrades the transthyretin messenger RNA transcript underlying hereditary transthyretin (hATTR) amyloidosis^2–4^. Within ON therapies as a class, microRNAs (miRNAs) offer perhaps the broadest range of therapeutic potential, ranging from clinical biomarkers to intervention strategies in cancer, fibrosis, hepatitis C, kidney and heart disease^5,6^. Importantly, microRNAs have been shown to regulate normal lung development, maintenance of different lung cell populations, and participate in the lungs response to injury and repair and their levels are changed in advanced lung disease, including Idiopathic Pulmonary Fibrosis (a chronic and lethal lung disease of unknown origin) ^7–12^. Among microRNAs, the miR-29 family has been extensively studied as a potential anti-fibrotic regulator based on its regulation of direct and downstream targets Collagen I and III, IGF1, and CTGF^13,14^. Although constitutively highly expressed, its expression levels are decreased in kidney^15^, lung^16,17^, liver, and myocardial fibrosis^18^, suggesting that supplementing miR-29 could be a therapeutic strategy for reversing or mitigating organ fibrosis. In earlier work, we demonstrated that a first-generation synthetic ON mimic of miR-29b; Remlarsen/MRG-201 (hereafter referred to as miR-201), blunted fibrosis in the bleomycin-induced pulmonary fibrosis mouse model^17^. In addition, in a randomized phase 1 clinical trial, intradermally administered miR-201 reduced collagen expression and delayed onset of fibroplasia in healthy volunteers, consistent with a broad anti-fibrotic therapeutic mechanism^19^. However, microRNA mimics are inherently unstable compounds and present numerous challenges for clinical translation^20^. Due to their size and charge, oligonucleotide-based compounds cannot passively enter cells, and are vulnerable to nuclease degradation^21^, sequestration in non-target tissues, and endo-lysosomal misrouting and degradation^1^. These vulnerabilities can be mitigated with backbone modifications (replacing a phosphate group with a sulfur atom) to create phosphorothioate (PS) linkages that increases circulation duration and hence reduce clearance via the kidneys^22^. In addition, ribose sugar modifications 2’-O-methylation or 2’-F-methylation helps limit nuclease degradation and increase plasma stability. Finally, therapeutic ON delivery can be increased and directed through covalent conjugation with bioactive molecules such as lipids, cholesterol (a general mechanism for boosting interactions with circulatory lipoprotein) or tissue-targeting agents such as peptides^23^.

Idiopathic Pulmonary Fibrosis (IPF) is a scarring, invariably lethal lung disease with two FDA approved therapies, which only slow down disease progression^24,25^. To realize the therapeutic potential of miR-29 mimicry in IPF we generated MRG-229, a next-gen miR-29 mimic with improved stability and potential for targeted delivery. In cultured human lung fibroblasts and lung slice cultures, we demonstrate that MRG-229 can reduce TGF-β induced fibrosis, as evidenced by downregulation of direct and downstream miR-29 targets *COL1A1* and *ACTA2*, respectively, at a 10-fold lower concentration than MRG-201. Similarly, in the bleomycin-induced mouse fibrosis model, we find that MRG-229 treatment counteracted upregulation of fibrosis-associated gene programs. At therapeutic dosing levels, MRG-229 therapy is associated with a favorable safety profile in mice, rats, and in non-human primates (NHPs). Finally, decreased concentrations of circulating mir-29 in the peripheral blood of patients with IPF are associated with substantially reduced survival. Taken together, these findings suggest that MRG-229 is an attractive preclinical candidate for therapeutic development in fibrosis-associated lung indications and potentially other fibrotic conditions.

## Results

### Design of the next-gen miR-29 mimic

MiRNA mimics are chemically synthesized double-stranded RNA molecules designed to elicit biologic activity by imitating mature miRNA duplexes. Such synthetic miRNA mimics must therefore be recognized by RNA-induced silencing complex (RISC) as an active compound. In contrast with miRNA inhibitors, for which only base pairing-mediated sequestration of its target mRNA is needed for activity, modification options of miRNA mimics are thought to be more constrained. Furthermore, to achieve therapeutic efficacy, such mimics must also be stabilized to ensure a longer half-live after administration^1^. To identify next-generation miR-29 mimics with improved *in vivo* stability, we performed an iterative discovery chemistry screen of MRG-201, our first-generation double strand miR-29 mimic^17^. To minimize hydrolytic degradation, we introduced additional 2’-O-methylation (2’OMe) or 2’-Fluoro (2’F) to the 2’ hydroxyl of nucleoside ribose moieties. Based on existing knowledge at the time MRG-201 was designed, we had introduced 2’F on bases in the active strand and 2’OMe on the bases in the delivery strand, that also has the conjugate. For our next-gen miR-29 mimic, we fully modified the parental MRG-201 compound with 2’F and 2’OMe as we removed all RNA from the molecule. In addition, we assessed bicyclic platelet-derived growth factor beta receptor (PDGFβR)-binding peptide (BiPPB), known to target cargo to pro-fibrotic cells for internalization^26^, for its potential to achieve targeted delivery. After having optimized the stabilization modifications, we conjugated BiPPB to our modified miR-29 mimic, which we named MRG-229 (Figure 1). We also conjugated the miR-29-mimic to a cholesterol moiety to be able to compare it to MRG-229 in downstream experiments.

**Figure 1:**
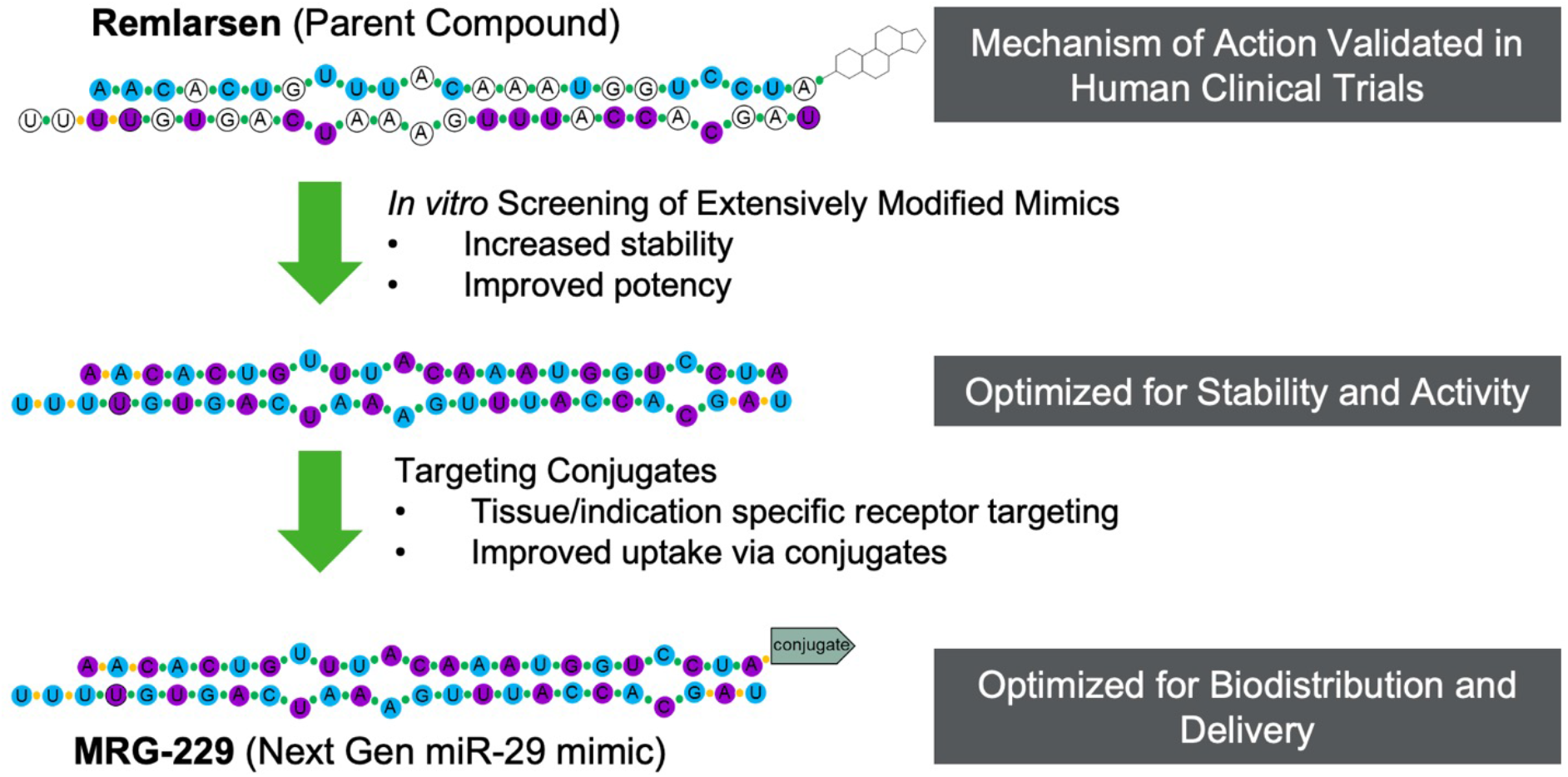
Overview of modifications differentiating second-gen MRG-229 from first gen MRG-201/Remlarsen. Top, first-gen MRG-201, the parent compound, bottom, MRG-229, the second-gen compound. DNA bases: white circles = unmodified base, blue circles = 2’OMe, purple circles = 2’F, linkages: green circles = phosphorodiester linkage, orange circles = phosphorothioate (PS) linkage. NH2 terminus modification: MRG-201/Remlarsen = cholesterol, MRG-229 = platelet-derived growth factor beta receptor (PDGFβR)-binding peptide (BiPPB),

### MRG-229 reduces fibrosis-associated phenotypes in cultured human lung fibroblasts and slice cultures

To confirm that the added modifications did not interfere with miRNA mimicking activity, we assessed MRG-229 in normal human lung fibroblasts (NHLFs) treated with TGF-β to induce pro-fibrotic genes. In control TGF-β-treated NHLFs, we found robustly increased expression of *COL1A1* (a direct miR-29 target). In contrast, in the presence of MRG-229, *COL1A1* and *ACTA2* gene expression levels were reduced in a dose-dependent manner (Figure 2A, B). These data demonstrate that MRG-229 retains the anti-fibrotic properties of the miR-29 mimic Remlarsen/MRG-201 through regulating the expression of miR-29 direct and downstream targets (Figure 2A-B). We next asked whether MRG-229-induced anti-fibrotic gene effects translated to regulation of Procollagen I C-peptide (PIP), a marker of newly synthesized, secreted collagen, without impacting on cell viability. To this end, we administered MRG-229 or MRG-201 to TGF-β treated NHLFs at concentrations ranging from 0.3-10 uM, assessed for cell viability using the WST-1 mitochondrial dehydrogenase assay, and collected supernatant samples in which we quantified PIP levels by ELISA. Cell viability in MRG-229-treated NHLFs was significantly improved relative to MRG-201-treated NHLFs, indicative of a higher tolerance of MRG-229. Although even low doses of MRG-201 reduced cell viability, with the highest 10uM dose being toxic (25% viability), only the highest dose of MRG-229 (10 μM) reduced viability (75%) with the second highest dose (5μM) being equivalent to control (Figure 2A). We also found that MRG-229 robustly down-regulated PIP levels, and at a superior potency relative to MRG-201 (Figure 2B). To assess whether similar effects could be achieved in already diseased cells, we treated fibroblasts from IPF patients (LL29 cells) with TGF-β followed by increasing concentrations of MRG-229. After 72 hours, we assessed overall collagen synthesis by assessing hydroxyproline levels by liquid chromatography-mass spectrometry. Relative to TGF-β treatment alone, MRG-229 significantly reduced cumulative hydroxyproline levels by 0.9 at 0.1μM, 1.1 at 0.5μM, 1.2 at 1 and 3μM, 0.7 at 10μM (Figure 2C). Together, these *in vitro* data demonstrate that MRG-229, with a greater potency and tolerability than Remlarsen, mitigates TGF-β-induced upregulation of fibrosis-associated genes and blunts collagen synthesis and secretion in normal and IPF lung cells.

**Figure 2:**
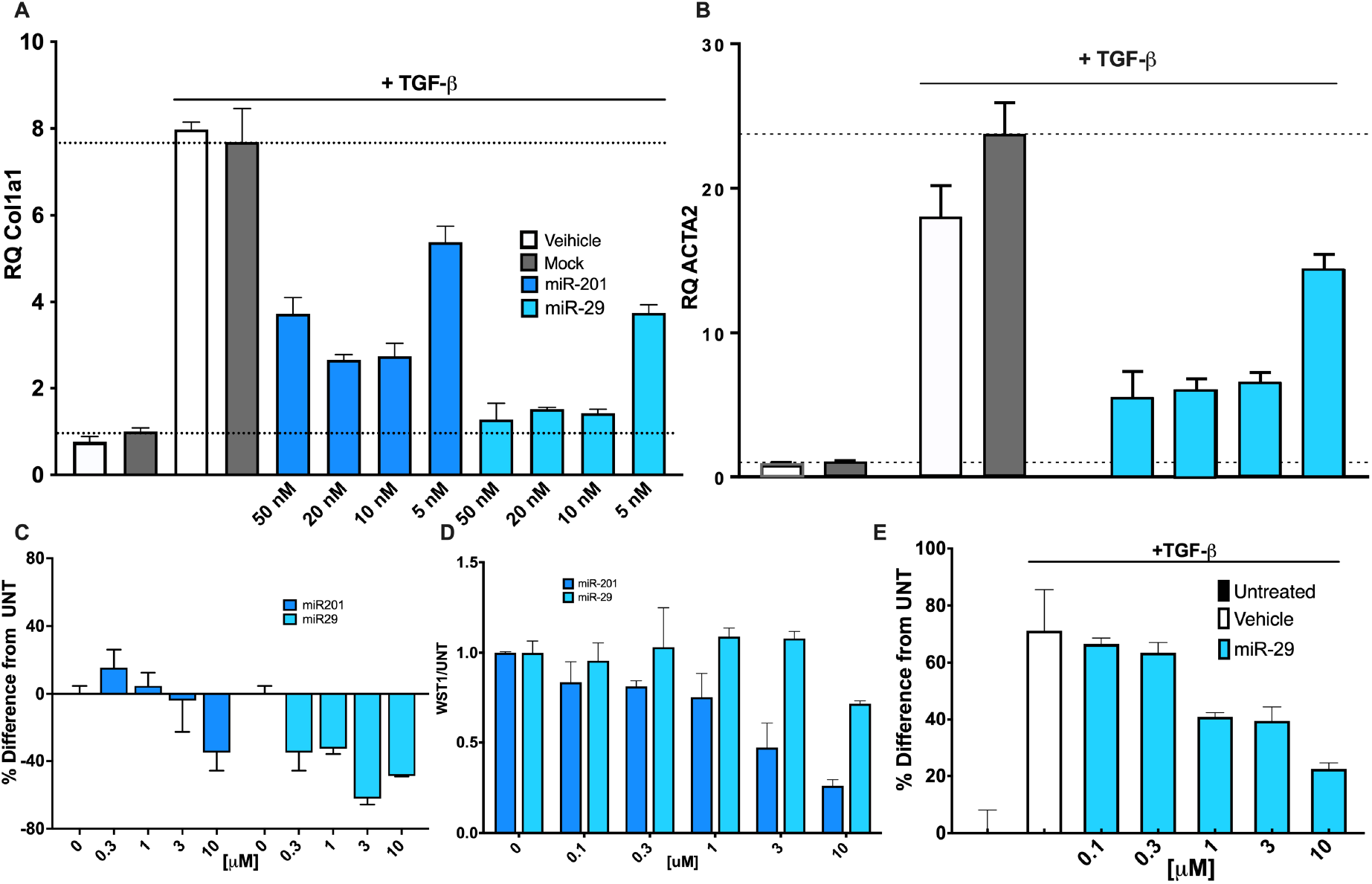
MRG-229 reverses fibrosis in vitro. (A) RT-PCR of the miR-29b target gene *COL1A* in NHLFs. (B) RT-PCR of the miR-29b target gene *ACTA2* in NHLFs. (C) Reduction of procollagen I C-peptide secretion in NHLFs treated with increasing concentrations of miR-201 (blue color columns) or miR-29 (light blue color column). (D) Cell viability in NHLFs treated with increasing concentrations of MRG-201 (THIS color columns) or MRG-229 (THIS color columns) (E) Dose-dependent collagen secretion assessed by accumulation of hydroxyproline levels in LL29 cells at 72h treated with TGF-Beta to induce fibrosis and treated with increasing concentrations of miR-29 as indicated. All experiments were performed on at least three biological replicates. Statistical analyses Ordinary one-way ANOVA were performed in GraphPad Prism.

Next, we sought to assess how MRG-229 regulates collagen production in the context of the normal physiological and architectural context of the human lung. To this end, we employed *ex vivo* culture of human Precision Cut Lung Slices (hPCLS), which can be cultured for up to 5 days, as previously described^27–29^. Briefly, cryopreserved 300μm thick sections of cadaveric human lung, derived from donors without a history of lung disease, were cultured and treated with control medium or medium containing a profibrotic cocktail (5ug TGF-b, 50ug PDGF-AB, 10ng TNFα, and 10mg LPA), after which we assessed *COL1A1* and *COL3A1* gene expression by qRT-PCR (figure 3A, B), and fibrosis-like phenotypes by collagen levels quantification after 5 days on histology (Figure 3C, D). When we treated hPCLS with MRG-229, we found that collagen production was significantly reduced, as assessed by quantitative imaging of histology after 120h of treatment (Figure 3A, B). As we had observed *in vitro*, MRG-229 significantly reduced expression of MRG-229 targets genes *COL1A1* and *COL3A1* in the *ex vivo* hPCLS model (Figure 3C, D). In both experiments, MRG-229 activity was comparable to the activity of the miR-29-mimic with a conjugated cholesterol moiety. The peptide conjugate and cholesterol conjugate to the non-targeting control sequence has no effect (Figure 3). Taken together, these data confirm that MRG-229 reduces experimentally induced fibrotic activity in both *in vitro* and *ex vivo* human disease models.

**Figure 3:**
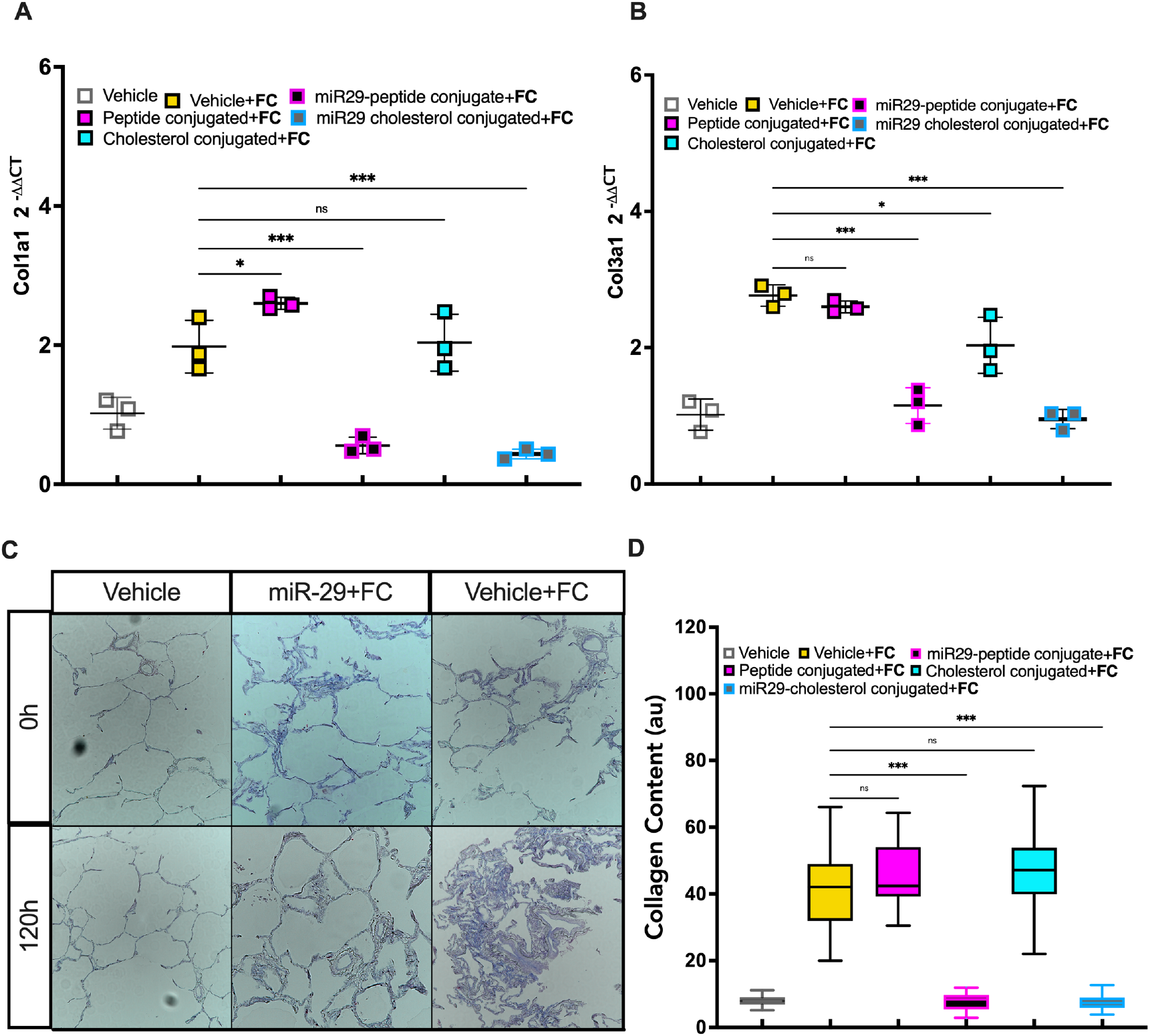
MRG-229 reverses fibrosis in a human Precision-Cut Lung Slice (hPCLS) model (blinded experiment). (A) RT-PCR analysis of COL1A1 relative expression (B) RT-PCR analysis of COL3A1 relative expression. (C) Masson Trichrome on human PCLS from healthy donors, cultured with either a fibrosis-inducing cocktail or vehicle for 120hrs with daily media changes, treated with miR-29 or a cholesterol-conjugated miR-29 mimic. (D) Graph depicts quantification of collagen levels after 5 days on histology. (n=3, Ordinary one-way ANOVA, *** = p<0.001)

### MRG-229 shows robust anti-fibrotic activity *in vivo*

To assess *in vivo* activity of MRG-229, we used two dosing paradigms in the bleomycin-induced pulmonary fibrosis mouse model. The first one was a prophylactic paradigm in which we administered compound at day 3 following bleomycin administration and collected tissue at day 14 (Figure 4A). First, we compared MRG-229 to our first generation miR-29 mimic MRG-201 in the prophylactic setting. Given that MRG-201 requires a 100 mg/kg dosing to achieve efficacy, we sought to assess whether we could lower the dosing for a next-generation compound by at least an order of magnitude, as that would be a starting point for a commercially viable dose. Accordingly, three days after bleomycin administration, we intravenously injected MRG-201 at 100 mg/kg and MRG-229 at 10 mg/kg twice weekly. In real-time PCR analysis of day 14 lung samples, we found a comparable down-regulation of miR-29 direct targets as well as non-direct targets (i.e., *CTGF*) in MRG-201 and MRG-229 treated bleomycin-injured mice (Figure 4C). In contrast, an unconjugated version of MRG-229 showed no *in vivo* activity compared to bleomycin/saline controls (Figure 4C). Similarly, bleomycin-treated animals injected with 100 mg/kg MRG-201 or 10 mg/kg MRG-229 showed reductions in total collagen content compared to bleomycin/saline controls as assessed by trichome staining and quantified as a percentage of total lung tissue (Figure 4D). Taken together, these dosing comparison experiments suggested that MRG-229 could be dosed at 10 mg/kg in mice to achieve a similar efficacy response as Remlarsen at 100 mg/kg.

**Figure 4:**
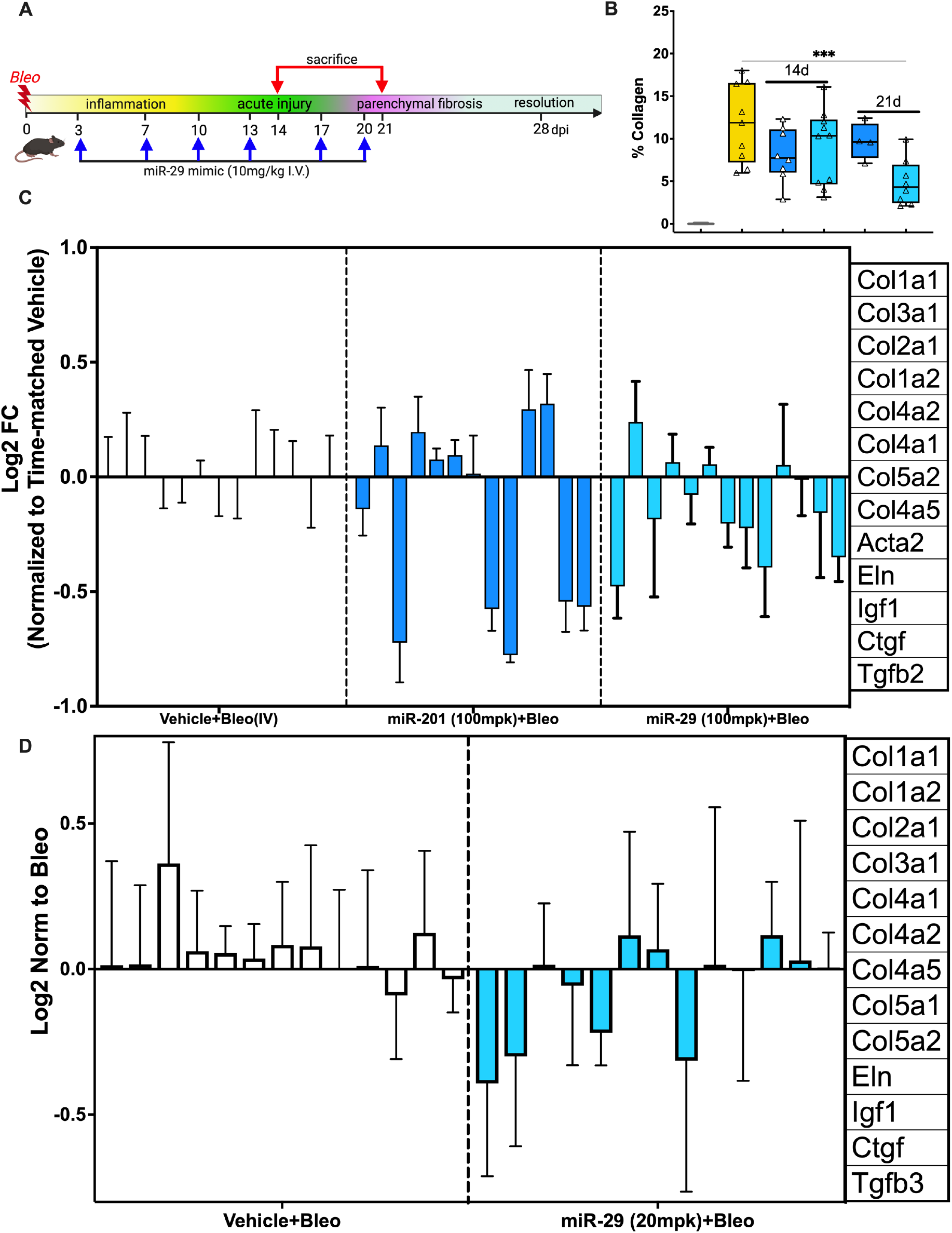
Day 14 analysis of prophylactic dosing of miR-201 at 100 mpk and miR-29 at 10 mpk in the bleomycin-induced lung fibrosis model. (A) Schematic of prophylactic dosing paradigm. Bleomycin-treated mice were dosed through intravenous injection with saline, **miR-201** at 100 mpk, or (n=3, Ordinary one-way ANOVA, *** = p<0.001). at 10 mpk, on days 3, 7, 10 and 13. On day 14, animals were sacrificed, and lung tissue analyzed. (B) Mean % total lung collagen quantified by Orbit machine learning image analysis software in bleomycin-induced mice treated with miR-201 or miR-29. (C) qPCR analysis of downregulated gene expression levels of a panel of fibrosis-associated genes in lung harvested from bleomycin-induced mice treated with either miR-201 or miR-29. C. (D) qPCR analysis of downregulated gene expression levels of a panel of fibrosis-associated genes in lung harvested from bleomycin-induced mice treated with either saline or the miR-29 mimic with all the stability modifications but lacking the BiPPB conjugate

We next asked if the anti-fibrotic effects observed with MRG-229 in mice extended to a therapeutic dosing paradigm (Figure 5A). Hence, we initiated twice-weekly dosing of 10 mg/kg MRG-229 10 days after bleomycin injury and collected tissue at day 21. In analysis of bronchoalveolar lavage fluid (BALF), we detected a ~20% reduction in IGF-1 levels, a known miR-29 target, and a ~40% reduction of TIMP1, a potential IPF biomarker in bleomycin-injured mice treated with MRG-229 relative to saline (Figure 5B, D). In quantitative histopathological analyses, we found that MRG-229 significantly reduced collagen deposition relative to saline, and preserved regions of normal alveoli architecture (Figure 5C). In addition, similarly to what we had observed for the prophylactic regimen, real-time PCR analysis of day 21 lung samples revealed down-regulation of miR-29 direct targets as well as non-direct targets in bleomycin-injured mice that received intravenous injection of MRG-229 (Figure 5E, right panel).

**Figure 5:**
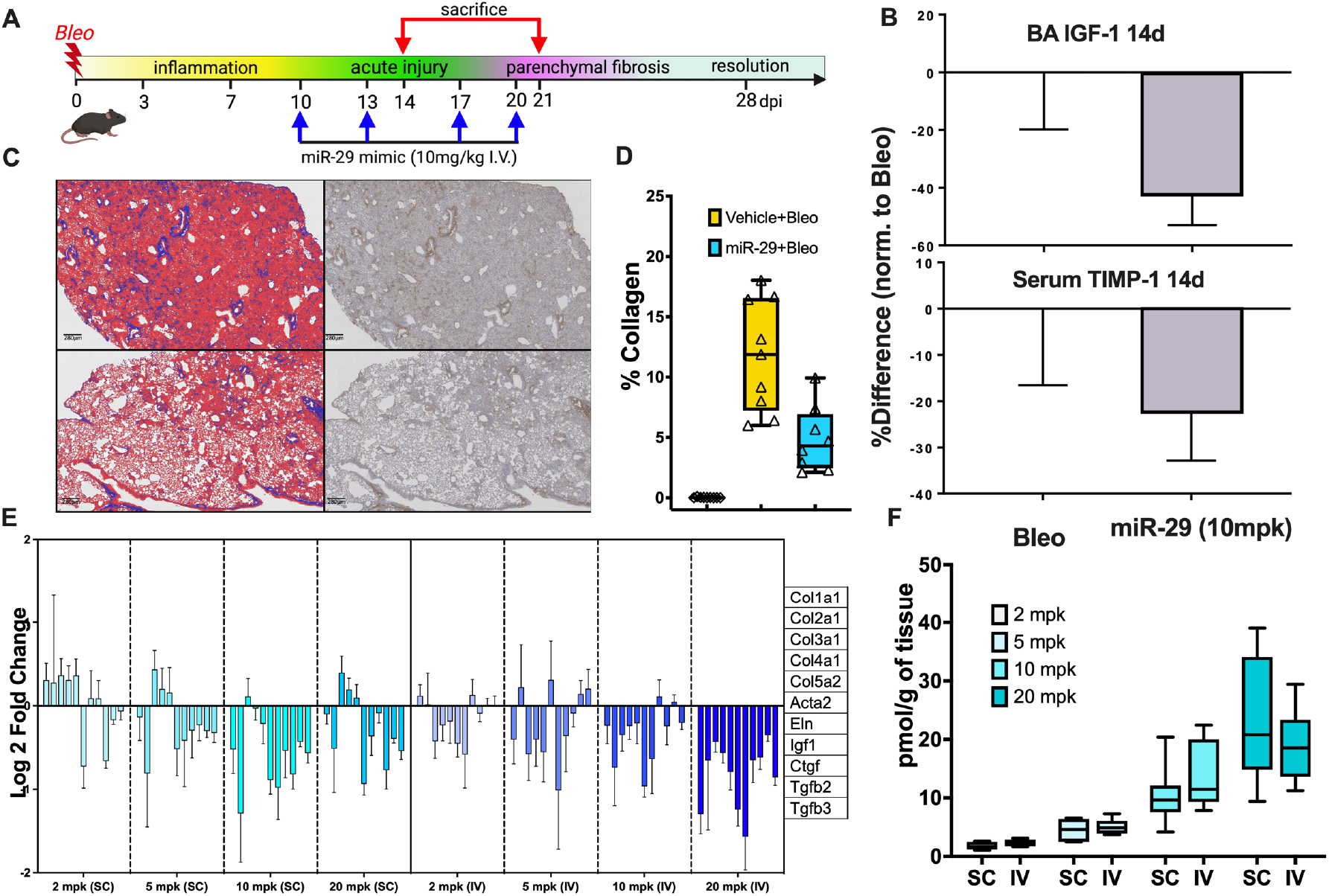
Day 21 analysis of therapeutic dosing of miR-201 at 100 mpk and miR-29 at 10 mpk in the bleomycin-induced lung fibrosis model. (A) Schematic of therapeutic dosing paradigm. Bleomycin-treated mice were dosed through intravenous injection with saline, miR-201 at 100 mpk, or miR-29 at 10 mpk, on days 10, 13, 17 and 20. On day 21, animals were sacrificed, and lung tissue analyzed. (D) Left: mean % total lung collagen quantified by Orbit machine learning image analysis software in bleomycin-induced mice treated with miR-201 or miR-29. (C) representative Masson Trichrome-stained histopathology images of saline-treated (top) and miR-201 -treated (bottom) lung sections. (B) ELISA analysis of IGF-1 levels in bronchoalveolar lavage fluid (left) and TIMP-1 levels in serum (right) harvested from mice treated with either saline and miR-29. D. Comparative analysis of subcutaneous and intravenous injection of miR-29 in bleomycin-induced mice, (E) qPCR analysis of downregulated gene expression levels of a panel of fibrosis-associated genes in lung harvested from bleomycin-induced mice treated with miR-29 as indicated. (F) lung distribution of miR-29 at different doses of either subcutaneous or intravenous injection.

Next, we asked whether subcutaneous administration of MRG-229 would achieve comparable *in vivo* efficacy to intravenous administration in the bleomycin-induced lung fibrosis model. Using regulation of pro-fibrotic genes as our readout for varying doses of MRG-229 at 2, 5, 10, or 20 mg/kg (administered either subcutaneously or intravenously) in the prophylactic paradigm, we found that subcutaneous MRG-229 dosing achieves therapeutic efficacy when administered at doses between 2 and 5 mg/kg (Figure 5E, left panel). When we next performed a sandwich-based ELISA assay from lung tissue homogenate to compare distribution in the lung upon intravenous and subcutaneous routes of administration, we found that MRG-229 distribution was comparable and dose-proportionate between subcutaneous and intravenous administration (Figure 5F). Taken together, these data support the potential for MRG-229 to be administered subcutaneously, while achieving therapeutic efficacy at a significantly lower and commercially viable dosing regimen than MRG-201.

### MRG-229 administration is associated with a favorable safety profile

Next, we comprehensively assessed MRG-229’s safety and toxicity profiles. From the doseresponse and route of administration studies in mice (Figures 4 and 5), our initial assessment of liver enzyme function and kidney damage markers showed that MRG-229 administration was not associated with any detrimental effect on either organ function (even in the presence of bleomycin) throughout up to 10 mg/kg biweekly dosing (Figure 6). Extending from these observations, we performed a 2-week repeat dose-range study of intravenous MRG-229 administration in Sprague Dawley Rats (Non-GLP). To this end, rats received formulation buffer (10 mM phosphate buffer diluted with isotonic buffered saline) or MRG-229 at 3, 10 or 30 mg/kg on Days 1, 4, 7, 11 and 14 (Table 1), after which we collected blood for hematology, coagulation, and serum chemistry analyses from the vena cava of fasted animals at necropsy. Using metabolic cages, we also collected and analyzed urine samples from fasted animals on Day 15. In addition, we performed gross pathology examinations and organ weight measurements on all animals at the terminal necropsy, and histopathology examination on all tissues from Groups 1 (Vehicle treated) and 4 (30 mg/kg MRG-229 treated), including gross lesions, liver, kidneys, spleen, lungs, and heart, from Groups 2 (3 mg/kg MRG-229 treated) and 3 (10 mg/kg MRG-229 treated) animals. Overall, we observed no measurable differences in body weight, food consumption or clinical observations (lethargy and grooming) between vehicle- and MRG-229-treated rats. Similarly, we did not observe any differences in hematology, clinical chemistry, coagulation, or urinalysis parameters. In histopathology analyses, we found that MRG-229 treatment was associated with minimal basophilic granularity in the tubular epithelium of the kidney with minimal tubular vacuolation found in one animal. In summary, MRG-229 administered intravenously at 3, 10, and 30 mg/kg twice weekly for two weeks in rats was well tolerated in both males and females, with no observable adverse effects at any dose tested.

**Figure 6:**
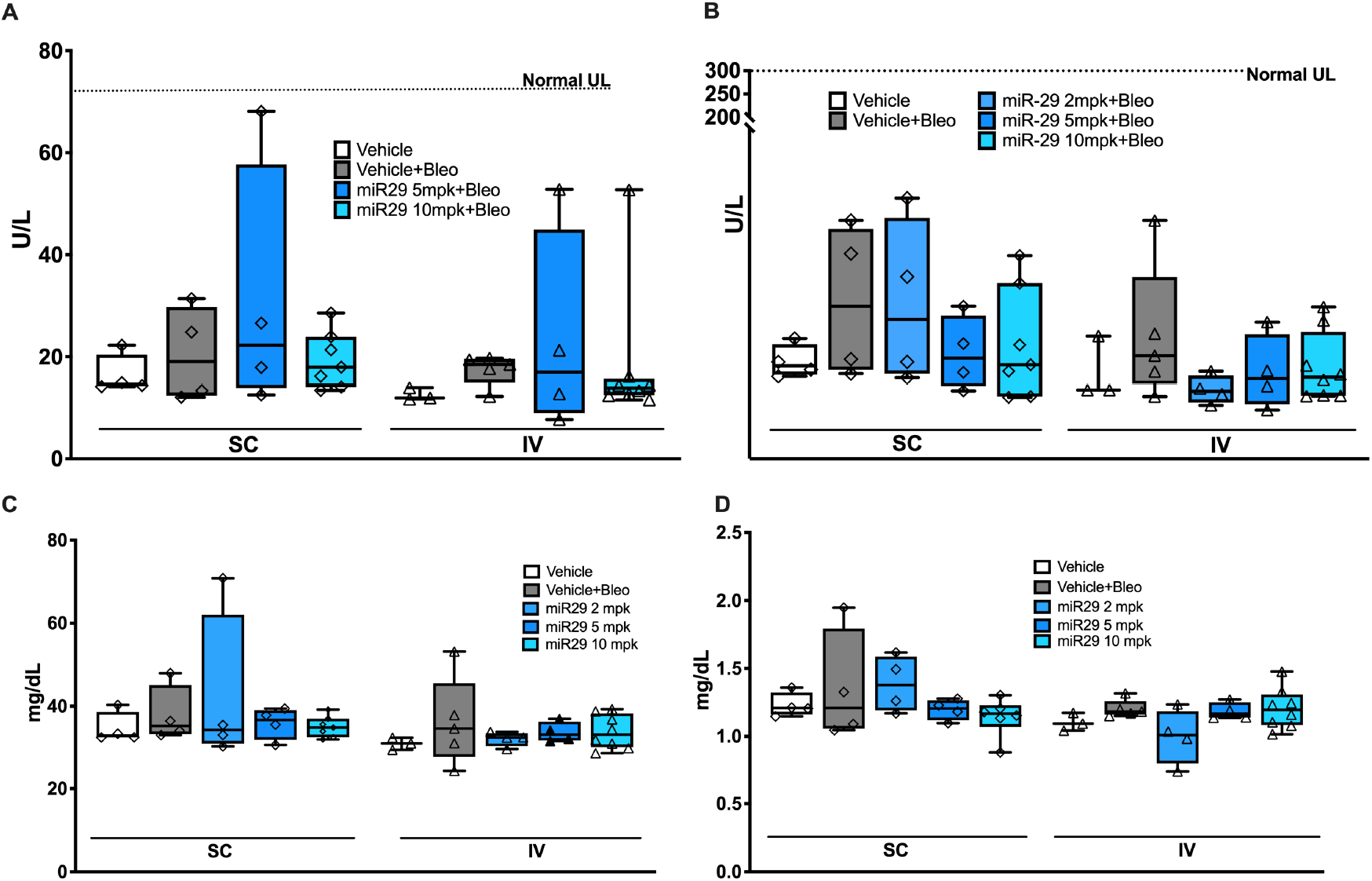
Liver (ALT and AST) and Kidney (Urea and Creatinine) assessments following MRG-229 administration. assessment of liver function enzymes and markers of kidney damage from the dose response and route of administration study showed no detrimental effect on liver or kidney function by MRG-229, even in the presence of bleomycin, up to 10 mg/kg biweekly administered

To assess potential toxicokinetic characteristics of MRG-229 by intravenous administration, we added three additional rats to Group 3 (10 mg/kg MRG-229) (Table 1). Samples for toxicokinetic analysis were collected on study Day 1 and study Day 15 before dosing and at 5 min, 30 min, 1 hr, 2 hrs, 4 hrs, 8 hrs, and 24 hrs after injection for all Group 3 animals. Animals in Groups 2 and 4 had toxicokinetic samples collected on study Day 1 and study Day 14 before dosing and at 5 minutes after injection. Plasma concentrations of MRG-229 decreased rapidly following the intravenous administration, with five out of six 24-hour plasma samples testing below the limit of quantification (BLOQ) (Supp Table 1; Supplementary Figure 1). Additionally, through concentrations (last dose, T=0) in all groups were BLOQ except for animal 2003 in the lowest dose group, which had a concentration of 5.8 ng/mL. We attribute this to analytical noise given that animal 3505 had a Day 1 pre-dose value with a similar result, even though animals had not yet been dosed with MRG-229. Select pharmacokinetic parameters for the TK animals were as expected (Supplemental Table 2). Last dose Cmax values across the three dose levels were approximately dose proportional, with a trend towards sub proportional increases in Cmax with increases in dose (Suppl Table 3) with no apparent gender differences observed (Suppl Table 4). Finally, to assess MRG-229 distribution to organ tissues, we collected samples from heart, liver, lung, kidney (including cortex), and spleen tissues from animals in terminal Groups 2-4 (Table 6). High concentrations of drug were detected in kidney tissue, with increases in concentration that were slightly lower than the increase in dose. Moderate amounts of drug were also measured in liver tissue, with less compound detected in spleen, heart, and lung tissues (Table 2).

### Intravenous delivery of MRG-229 in non-human primates shows no adverse effects

Toxicological studies of potential therapeutic compounds in non-human primates are critical to assess their potential for translation to the clinic. To this end, we assessed MRG-229 in a dose range finding study in non-human primates (NHPs). We administered MRG-229 by IV injection to naïve cynomolgus monkeys (1 animal/sex/group) at concentrations of 0, 5, 15, and 45 mg/kg on Days 1, 4, 7, 11, and 15, and performed necropsy and sample collection on Day 16. We did not observe any notable parameters related to clinical observations, food consumption, and bodyweights. Similarly, we found no evidence of MRG-229 related findings in the hematology, clinical chemistry, coagulation, or urinalysis parameters assessment. In all dose groups, including vehicle, we found evidence of a slight decrease in hematocrit and an increase in reticulocytes on Day 2 and Day 16 relative from pretreatment values, likely due to the extensive blood sampling protocol requirements. We observed increased AST and CK values on Day 2 but these had generally resolved by Day 16 in all treatment groups, including the female vehicle control animal. Due to the lack of a clear dose response relationship, and the fact that we observed the similar changes in the control animals, these findings are unlikely to be related to MRG-229. We did not identify any MRG-229-related histologic findings in tissues from Group 4 (45 mg/kg MRG-229) animals (Suppl Table 5), suggesting that the dose tested is safe.

Following IV administration, MRG-229 plasma concentrations decreased rapidly (and by 24 hours, concentration of MRG-229 had decreased to less than 0.5% of initial values, with several samples testing below the quantifiable level (BQL) (Suppl Table 6). Select pharmacokinetic parameters are reported in Suppl Table 7, showing individual animal and mean results (no other statistics are reported due to the small samples sizes). There was no apparent difference between male and female animals, with similar PK results for both animals in all groups. As expected for IV bolus, Tmax was at the time of the first sample collected after dosing (5 minutes) for almost all animals. Cmax concentrations were expectedly high following the intravenous bolus dose, reaching Day 1 mean values of 312 μg/mL for the MRG-229 45 mg/kg group. There was no accumulation observed for either Cmax or AUClast values from Day 1 to Day 15 (data not shown), however Cmax and AUClast values across the MRG-229 dose groups were nearly perfectly proportional, with dose normalized values being very similar across dose group for both Day 1 and Day 15 PK curves (Suppl Table 7), and nearly perfectly linear increases in Cmax and AUClast values across the dose range were observed (Suppl Table 8). Finally, distribution results to organ tissues 24 hours after the final dose are listed in Table 3. As expected, MRG 229 was detected in the lung, as well as in other tissues, consistent with our earlier mouse data.

### Exosomal miR-29b levels in plasma and serum predict mortality in a cohort of IPF patients

Given that delivery of a miR-29 mimic reduces fibrosis in animal models, we reasoned that miR-29 levels in humans could be used to develop a precision medicine-based approach to identify individuals with low miR-29 levels at risk of death. To potentially identify such patients, we examined circulating and exosomal miR-29b levels in a cohort of 46 and 213 patients with IPF diagnosis recruited from Yale and Nottingham Universities (Profile Cohort), respectively. Table 1 shows the clinical characteristics of IPF patients in both cohorts. For each patient, we performed miRNA extraction from plasma and serum samples isolated from blood, followed by multiplexed, color-coded probe pairs to assess levels of miR-29b specifically. miRNA data was normalized using top 100 normalization and log2 transformed miR-29b levels were used for statistical analysis. Receiving Operating characteristics (ROC) curves were used to determine the optimal threshold for mortality prediction using exosomal miR-29b in IPF patients from both Yale and Profile cohorts. Cox proportional hazard’s models and Kaplan-Meir curves were used to determine the association between exosomal miR-29b levels, adjusted to GAP index, and IPF mortality.

We found that ROC identified similar miR-29b exosomal RNA thresholds for mortality prediction in the Yale (plasma level threshold of 4.84) and Profile (serum level threshold of 4.32) cohorts. After adjusting for the GAP severity index, exosomal miR-29b levels in plasma (≤ 4.86) and serum (≤ 4.32) were significantly predictive of mortality in the Yale (HR:0.156, 95%CI: 0.0404-0.6066, P=0.0073) and Profile (HR:0.5066, 95%CI: 0.2984-0.8599, P=0.011) cohorts, respectively (Figures 7A, B).

**Figure 7.**
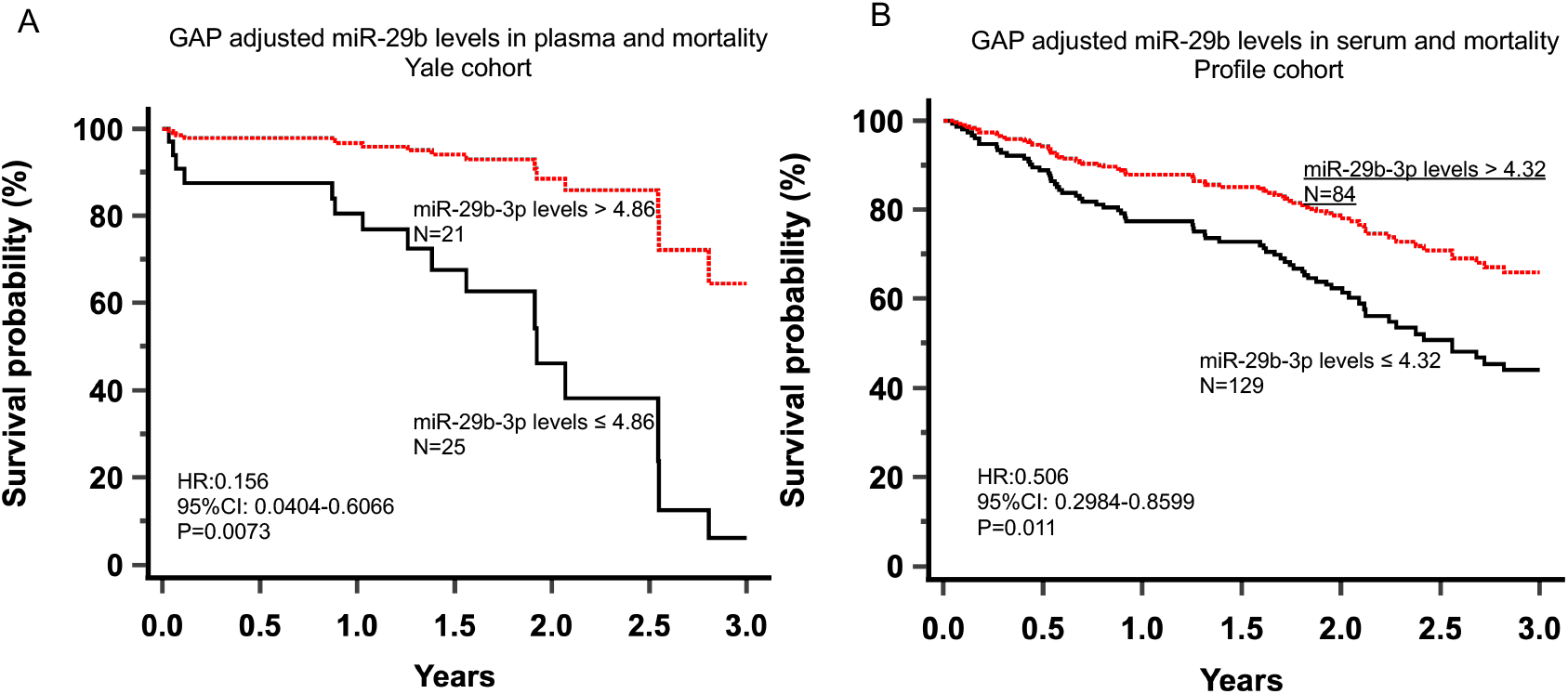
miR-29 survival in 2 IPF cohort: ROC identified similar miR-29b exosomal RNA thresholds for mortality prediction in the Yale (plasma level threshold of 4.84) and Profile (serum level threshold of 4.32) cohorts. After adjusting for the GAP severity index, exosomal miR-29b levels in plasma (≤ 4.86) and serum (≤ 4.32) were significantly predictive of mortality in the Yale (HR:0.156, 95%CI: 0.0404-0.6066, P=0.0073) and Profile (HR:0.5066, 95%CI: 0.2984-0.8599, P=0.011) cohorts, respectively. Figures 7A and 7B.

## Discussion

In this study, we report the generation of MRG-229, a next-gen miR-29 mimic capable of reversing fibrosis-associated molecular transcription and secretion phenotypes in human cellular lung fibrosis models. Importantly, we found that MRG-229 is effective and safe in the lungs at low doses. Relative to our first-gen compound MRG-201, MRG-229 achieved comparable levels of fibrosis reversal *in vitro* at a ten-fold lower dosing level, an improvement which enabled us to explore both efficacy and safety parameters using commercially viable doses in preclinical animal models. In bleomycin-induced mice, we found that MRG-229 effectively achieved downregulation of direct and indirect miR-29 profibrotic target genes concomitant with reduced collagen secretion and preserved lung alveolar architecture. We also found that MRG-229 administration in mice was associated with a favorable safety profile at 10 mg/kg dosing. Based on these data, we increased the maximum dose in a two-week repeat-dosing study in rats to 30 mg/kg (5 dosing events over each two week-period) and obtained equivalent safety results. In our NHP toxicology analysis, we increased the highest dose to 45 mg/kg (5 dosing events over 15 days) and did not detect any clinical pathology alterations at the highest dose that could be attributed to IV administration of MRG-229. Intravenous administration is inconvenient to patients and associated with a higher risk for adverse events relative to oral and subcutaneous delivery routes. In bleomycin-induced mice, we assessed both intravenous and subcutaneous MRG-229 administration and found that both delivery approaches reduced pro-fibrotic gene expression programs at therapeutically relevant doses. We further demonstrated that levels of miR-29 in both the serum and plasma may be predictive of mortality in IPF patients and could be used to identify patients that could have a survival benefit from MRG-229 administration. Taken together, these data suggest that administration of MRG-229 is safe and effective at commercially viable dosing levels and may be an attractive candidate for treatment of IPF.

It is well established that miR-29 family members can regulate extracellular matrix proteins and achieve therapeutic effects when elevated in organs susceptible to fibrosis, including the lung^13,30^. A recent study reported the results of a Ph1 double-blinded, placebo-controlled clinical trials in healthy volunteers (n=47) that did or did not receive skin incisions, and that were treated with intradermal miR29b mimetic MRG-201 or placebo injections. In this trial, MRG-201 had no impact on normal wound healing but significantly decreased fibroplasia relative to placebo. This study serves as a proof of concept for MRG-201 therapy in human skin as an approach to prevent formation of a fibrotic scar^19^. In the context of our paper, this is an important result as it demonstrates that miR-29 has an antifibrotic effect *in-vivo* in humans. We have shown that delivery of MRG-201 blocked pulmonary fibrosis in a mouse model^17^; however, this was achieved at dosages that would never be therapeutically acceptable in humans. To overcome this limitation in a compound that was otherwise effective we generated MRG-229, a compound with 2’F and 2’OMe modifications and conjugated to the bicyclic platelet-derived growth factor beta receptor (PDGFβR)-binding peptide (BiPPB) known to target pro-fibrotic cells for internalization^26^. MRG-229 proved to have the same antifibrotic effects as MRG-201, but with distinct pharmacological advantages, as it more stable in-vivo, can be administered in low doses and subcutaneously with good detection in the lung. To this we now add the evidence that MRG-229 has antifibrotic effects in the human lung ex-vivo using PCLS, an excellent experimental paradigm to investigate human lung biology as it preserves much of the cellular and molecular functions of lung tissue. Furthermore, each slice of lung tissue is 300um thick, which also allows preservation of the spatial complexity. MRG-229 led to reversal of fibrosis in human PCLS treated with a profibrotic cocktail, an accepted ex-vivo translational model of pulmonary fibrosis. Considering that we have seen an antifibrotic effect in-vivo in humans (in the skin) with MRG-201, we feel that our PCLS results establish a strong case that miR-29 mimicry will be antifibrotic in humans, and that MRG-229, given its safety profile, stability and superior pharmacodynamic properties, may be a suitable agent to confer this effect.

A common challenge with IPF therapeutics is that there is little evidence that the target mechanism is in fact implicated in the patients we treat. This is of course even more complex with the two FDA approved drugs, in which data about mode of action is limited. In the case miR-29, the therapeutic premise is relatively straightforward: miR-29 is reduced in nearly every fibrotic condition, and in the lung, it has been shown repeatedly to be decreased^13,31,32^. McDonough et al assessed gene expression changes in the differentially affected regions in the IPF lung^32^, and demonstrated that miR-29 was decreased in the IPF lung even in relatively conserved areas, whereas genes known to be regulated by miR-29 were increased as fibrosis progressed. Unfortunately, it is impossible to assess miR-29 in the lung in most patients as biopsies are limited. In this manuscript, we provide a novel observation that may be very important in this context. We demonstrate that in two independent cohorts decreased peripheral blood exosomal miR-29 is associated with worse prognosis in patients with IPF. Though it may be too early to determine whether treatment with miR-29 mimics should be offered preferentially to patients with low blood levels, nonetheless this finding highlights the connection between reduced miR-29 levels and IPF progression, and potentially could be used in the future as a companion diagnostic.

In this study, we did not investigate the biological mechanism underlying the action of miR-29 mimic, as this was established in our previous work in the mouse lung^17^. Mouse and human lung share a highly similar cellular composition, function and architecture and pathways of gene regulation are broadly conserved between mouse and human. As in mouse, we observed reduced levels of Col1a1 and Col3a1 in the presence of miR-29 mimic in the bleomycin-treated mice, consistent with the known role of miR-29 in regulating many different extracellular matrix related genes^33^. At the cellular level, it would be interesting to know which cell type/s in the human PCLS lung tissue are the primary mediators of the observed antifibrotic response. It would also be interesting (though difficult) to address which cell types are primarily responsible for declining miR-29b levels in the peripheral blood from the two IPF cohorts, as this may represent an attractive target for cell therapies. In our study, levels of MRG-229 were detected not only in the lung but also in several other organs. Such systemic delivery is an important limitation of oligo-based therapeutics in general; however previous studies have demonstrated that there is minimal on-target activity or pharmacodynamics in tissues where the miRNA is not dysregulated. Longer toxicology studies will need to be performed prior to more chronic dosing in humans, while inhalation as an alternative mode of administration may help to increase targeting of MRG-229 specifically to the lung.

IPF affects more than 5 million people worldwide and has a poor prognosis^2,24,34–36^. Despite the heavy toll of IPF, only two FDA-approved therapies exist for IPF, neither of which reverse fibrosis or improve the well-being of treated patient. Due to the breadth and complexity of aberrations in many signaling, aging, development, and stress pathways in IPF, a broad-acting therapeutic mechanism based on miRNAs that can target multiple pathways represents an attractive therapeutic candidate. Indeed, the IPF lung exhibits significantly altered patterns of microRNAs expression ^7–9,12,37^ and low blood plasma and serum levels of miR29-b are predictive of mortality in IPF patients supports the use of miR-229 as a potential therapy. Taken together, the findings reported here represent a solid range of IND-enabling work for further clinical development of MRG-229 in pulmonary fibrosis indications.

## MATERIALS AND METHODS

### Animals

All animal studies were conducted in accordance with the NIH guidelines for humane treatment of animals and were approved by the Animal Care and Use Committee (IACUC) at MiRagen Therapeutics, Inc. and at Yale University.

### Cell culture

Normal Human Lung Fibroblasts (NHLFs), obtained from Lonza, were cultured in Fibroblast Basal Medium (FBM) supplemented with Fibroblast Growth Factor (FGF) and maintained at 37°C and 5% CO_2_. LL 29 (ATCC CCL-134™) cells were maintained in Ham’s F12K medium with 15% FBS and maintained at 37°C and 5% CO_2_. Proliferation and viability was assessed using the WST-1 assay (Sigma).

### Oligonucleotide synthesis

*In vitro* and *in vivo* studies utilized oligonucleotides that were produced via solid phase support synthesis at miRagen Therapeutics, Inc. (Boulder, CO) and formulated in PBS. All oligonucleotides were sterile filtered during formulation. Vehicle control for animal studies was sterile PBS.

### Quantitative Real-time PCR Analysis

To quantify *in vitro* regulation of pro-fibrotic genes *COL1A1* and *ACTA2*, NHLFs were treated with 5ng/mL TGF-b (PeproTech) to induce the fibrotic response. Cells were then treated with MRG-229 through passive administration at concentrations ranging from 0.3 to 10 mM or through transfection at concentrations ranging from 5nM to 50nM at 0.2 mL/well Dharmafect I (Thermofisher Scientific) as per the manufacturer’s instructions. After 72h, samples were harvested and mRNA expression levels analyzed with qPCR (Life Technologies) using human Col1a1 and Acta2 primers from ABI for the specified gene and species.

For *in vivo* real-time PCR analysis, 50-100 mg of tissue was homogenized in 1 mL Trizol (Invitrogen) in Lysing Matrix D tubes (MP Biomedical) with shaking for 4 × 20 seconds using an Omni BeadRuptor 24 (Omni Intl.) at a speed of 5.65. Total RNA was extracted as per the manufacturer’s standard protocol, after which 5 μL of 125 ng/μL RNA from each tissue sample was used to generate cDNA using Applied Biosystems High Capacity cDNA Reverse Transcription Kit per manufacturer’s specifications. The expression of a subset of genes was analyzed by quantitative real time PCR using Taqman probes purchased from Thermo Fisher Scientific.

### Hydroxyproline assay in cultured cells

Samples of untreated and TGF-b treated LL 29 cells, and samples that had also received increasing concentrations of MRG-229, were dried until constant weight and hydrolyzed in 12 N HCl for 3 h at 120°C. Hydroxyproline was then detected using an in-house LC MS/MS assay developed at MiRagen. Data are expressed as percent difference relative to untreated LL 29 cells.

### Procollagen IC-peptide (PIP) analysis

Secretion in PIP in NHLFs treated with MRG-201 or MRG-229 was conducted by collecting the supernatant of treated cells and performing procollagen type I PIP ELISA (Takara).

### Human precision cut lung slice (hPCLS) culture

Human lung segments were obtained through the National Disease Research Interchange (NDRI) and hPCLS generated as previously described^29^. Briefly, low melting-grade agarose (3 wt-%) was slowly injected via a visible bronchus to artificially inflate the lung segments. Segments were cooled at 4°C for 30 minutes to allow gelling of the agarose and then cut to a thickness of 300μm using a Compresstome (VF-300-0Z by Precisionary) at cutting speed of 6 μm s^−1^ and oscillation frequency of 5Hz. The hPCLS were cultured in 24 multiwell plates (Corning) in 500μL DMEM-F12 no-phenol red containing 0.1% FBS and 1% penicillin/streptomycin.

### Fibrosis model in hPCLS cultures

hPCLSs, 3 slices for each experimental condition, were exposed to either a fibrosis-inducing (including TGF-Beta; 5ng/ml, TNF-alpha; 10 ng/ml, PDGF-AB; 5μM, and LPA; 5μM) or a control cocktail (including vehicle control) for 120h with media replaced every 24h as previously described^29^. In this blinded experiment, fibrosis-induced slices were treated with MRG-229 (sample #2), a cholesterol-conjugated miR-29 mimic (sample #4), each at 200μM final concentration, or control (samples #1, #3). After 5 days after treatment, lung slices were processed for analysis; divided for histology and RT-qPCR analysis, respectively.

### Histology on hPCLS

hPCLS were fixed with 4% (weight/volume) paraformaldehyde overnight, and paraffin-embedded at 0h and 120h. 3μm sections were cut using a microtome, mounted on glass slides and subjected to antigen retrieval. After deparaffinization and rehydration, staining was performed according to standard protocols for Masson’s Trichrome, and samples mounted using mounting medium and covered with a cover slip. For collagen quantification, bright field scanning with a Nikon inverted microscope at 20X magnification was used to acquire two representative images for each sample and at least 20 different random field of views. Collagen staining was quantified and determined by percentage of stained areas. Images were analyzed with ImageJ software (ImageJ NIH).

### Histology

Samples were sent to HistoTox Labs for paraffin embedding, sectioning, and staining with Hematoxylin and Eosin and Masson’s Trichrome. Masson’s Trichrome sections were evaluated for collagen deposition and fibrotic mass using Orbit image analysis. Full sections were evaluated for the % collagen in the whole tissue, for the % collagen in the lung tissue and fibrotic mass (excluding structural collagen), for the % collagen in the lung tissue (excluding structural collagen and fibrotic mass), and for the % tissue area that was fibrotic mass (excluding structural collagen and normal lung tissue).

### Biodistribution Analysis

A sandwich hybridization assay was used for the quantification of promiR-29 in tissue samples. Probes for the hybridization assay were synthesized using 2’Ome, and LNA modified nucleotides. Detection was accomplished using anti-fluorescence-POD, Fab fragments (Roche) and TMB Peroxidase Substrate (KPL). Standard curves were generated using non-linear logistic regression analysis with 4 parameters (4-PL). The working concentration range of the assay was 2-2000 ng/mL. Tissue samples were prepared at 100 mg/mL by homogenizing in 3M GITC buffer (3 M guanidine isothiocyanate, 0.5 M NaCl, 0.1 M Tris pH 7.5, 10 mM EDTA) for 2 × 45 seconds using an Omni BeadRuptor 24 (Omni Intl.) at a speed of 5.65. Tissue homogenates were diluted a minimum of 50-fold in 1 M GITC Buffer (1 M guanidine isothiocyanate, 0.5 M NaCl, 0.1 M Tris pH 7.5, 10 mM EDTA) for testing.

### RNA isolation and RT-qPCR on PCLS

hPCLS samples were snap frozen in liquid nitrogen and homogenized using a hand-held homogenizer. QIAGEN miRNA was used for total RNA isolation. The RNA concentration and quality were assessed using NanoDrop spectrophotometer (Thermo Fisher Scientific, Germany). Relative expression of MiR-29 targets genes COL1A1 and COL3A1 mRNA levels from all *ex vivo* experiments were determined by RT-qPCR on Viia7 1.0 Real-Time PCR system using TaqMan gene expression assays. Reverse transcription with random primers and subsequent PCR were performed with TaqMan RNA-to-CT 1-Step Kit (Applied Biosystems). All experimental groups were assessed as 6 technical replicates and repeated at least three times.

Raw data for cycle threshold (Ct) values were calculated using the ViiA7 v.1 software with automatically set baseline. The results were analyzed by the ΔΔCt method and normalized to GAPDH. Fold change was calculated by taking the average over all the control samples as the baseline. All the probes used in this study were purchased from Thermo Fisher Scientific.

### Bleomycin model of pulmonary fibrosis

Male C57Bl/6 mice, 9-10 weeks of age, were purchased from Taconic Biosciences, Hudson, NY and allowed to acclimate for at least on week prior to experiments. Mice were anesthetized with dexmedetomidine, 1 mg/kg IP, intubated, and given one, intra-tracheal dose of bleomycin at 1.25 mg/kg in 50uL saline or an equivalent volume of saline. A reversal agent was given subcutaneously once the mice had been dosed. Animals were then treated with MRG-201, MRG-229, or an equivalent volume of 0.9% saline by intravenous injection and euthanized on days 8, 14, and 21 after bleomycin administration. Serum and BALF samples were collected as well as tissue samples. Serum was collected and used for measurement of ALT, AST, BUN, and creatinine activity. Bronchoalveolar lavage fluid (BALF) was collected, spun at 1200 × g for 15 min after which supernatant and pellet were separated and frozen. BALF supernatant will be decanted/collected and placed in a separate 1.5 mL Eppendorf tube. BAL cell pellets and supernatant will then be frozen and stored. The left lobe of the lung was dissected and used for histology and molecular assessment, whereas the right caudal lobe was flash frozen in liquid nitrogen and used for hydroxyproline/collagen assays and biodistribution analysis. Liver, kidney, spleen, and heart tissue was also collected and flash frozen.

### Exosomal miRNA isolation from IPF cohorts

#### Participants

46 and 213 patients with IPF diagnosis were recruited from Yale and Nottingham Universities (Profile Cohort), respectively. IPF diagnosis was based on guidelines of the American Thoracic Society and European Respiratory Society (ref). Patients were followed until death or loss of follow up. Follow up time was limited to three years.

##### Sample collection, Yale University cohort

Blood was collected in heparin tubes using a routine procedure and was immediately (within 10 minutes after blood collection) centrifuged at 4°C, 1200×g, for 10 min). Plasma was aliquoted and frozen at −80°C until analysis. All patients provided written informed consent and a protocol incorporating biomarker-studies was approved by the Institutional Review Board (IRB), Yale School of Medicine (HIC#0706002766).

##### Sample collection, Profile cohort

###### Exosomal miRNA isolation

Briefly, 400μl plasma or serum were used as starting material for exosomal miRNA’s extraction using the Norgen Biotek Plasma/Serum Circulating and Exosomal RNA Purification Mini Kit (Thorold, Ontario, Canada). To detect circulating and exosomal miR-29b levels, we used multiplexed, color-coded probe pairs (Nanostring nCounter analysis system) using 20 ng of total RNA, following the manufacturer’s protocol.

###### Statistical analysis

miRNA data was normalized using top 100 normalization and log2 transformed miR-29b levels were used for statistical analysis. Receiving Operating characteristics (ROC) curves were used to determine the optimal threshold for mortality prediction using exosomal miR-29b in IPF patients from both Yale and Profile cohorts. Cox proportional hazard’s models and Kaplan-Meir curves were used to determine the association between exosomal miR-29b levels, adjusted to GAP index, and IPF mortality.

## Supplementary Materials

Table S1. Clinical profile characteristics of IPF patients in both cohorts

Table S2. Experimental Study Groups and Dose Information

Table S3. Distribution of MRG-229 Antisense strand to organ tissues in rats

Table S4. Distribution of MRG-229 Antisense strand to organ tissues in NHPs

Table S5. Plasma Concentrations of MRG-229 Antisense Strand in Rats

Table S6. Pharmacokinetic Parameters of TK Animals in Rats

Table S7. Dose Proportionality of Last Dose Cmax Values in Rats

Table S8. Male vs Female Cmax Values in Terminal Animal Groups

Table S9. Plasma Concentrations of MRG-229 Antisense Strand in NHPs

Table S10. Mean and Individual Pharmacokinetic Parameters in NHPs

Table S11. C_max_ and AUC_last_ Dose Proportionality in NHPs

## Author contributions

NK, RLM, MC, GY, SR, KR, and LP conceived, designed, and analyzed experiments and results. MC, KR, RN, GY, NA, FA, JHM, and OD performed and analyzed results. MC, RLM and NK wrote the manuscript.

## Data and materials availability

All data are available in the main text or the supplementary materials.

