## Supplementary material for "A lung targeted miR-29 Mimic as a Therapy for Pulmonary Fibrosis": Supp Table 1

**Supplement**

**Supp Table 1: Clinical profile characteristics of IPF patients in both cohorts.**

|  | Yale cohort (N=46) | Profile cohort (N=213) |
| --- | --- | --- |
| Age (± SD) | 68.5 (± 7.5) | 73 (± 6.62) |
| Gender |  |  |
| Male (%) | 39 | 166 |
| Female (%) | 7 | 47 |
| FVC% (± SD) | 75.77 (± 9) | 80.26 (± 22.15) |
| DLCO% (± SD) | 49.1 (± 9.6) | 41.7 (± 21.5) |
| GAP index (± SD) | 3.96 (± 1.41) | 2.56 (± 0.52) |

Supp Table 2. Experimental Study Groups and Dose Information

|  |  |  | **Dose Level (mg/kg)** | **Dose Concentration (mg/mL)** | **Dose Volume (mL/kg)** | **Terminal** | | **TK** |
| --- | --- | --- | --- | --- | --- | --- | --- | --- |
| **Group** | **Species** | **Test Material** |  |  |  | **M** | **F** | **F** |
| 1 | Rat | Control | 0 | 0 | 2 | 3 | 3 | 0 |
| 2 | Rat | MRG-229 | 3 | 1.5 | 2 | 3 | 3 | 0 |
| 3 | Rat | MRG-229 | 10 | 5 | 2 | 3 | 3 | 3 |
| 4 | Rat | MRG-229 | 30 | 15 | 2 | 3 | 3 | 0 |
| 1 | NHP | Control | 0 | 0 | 1 | 1 | 1 |  |
| 2 | NHP | MRG-229 | 5 | 5 | 1 | 1 | 1 |  |
| 3 | NHP | MRG-229 | 15 | 15 | 1 | 1 | 1 |  |
| 4 | NHP | MRG-229 | 45 | 45 | 1 | 1 | 1 |  |

Supp Table 3: Distribution of MRG-229 Antisense strand to organ tissues in rats

| **Group and Dose** | **Animal # F/M** | **Concentration (µg/g)** | | | | | | | | | |
| --- | --- | --- | --- | --- | --- | --- | --- | --- | --- | --- | --- |
|  |  | **Kidney** | | **Liver** | | **Spleen** | | **Heart** | | **Lung** | |
|  |  | **Female** | **Male** | **Female** | **Male** | **Female** | **Male** | **Female** | **Male** | **Female** | **Male** |
| Group 2 | 2501 / 2001 | 80.3 | 91.2 | BLOQ | 0.248 | BLOQ | 0.247 | BLOQ | BLOQ | BLOQ | 0.525 |
| 3 mg/kg | 2502 / 2002 | 80.3 | 53.2 | BLOQ | BLOQ | 0.840 | BLOQ | BLOQ | 0.154 | BLOQ | MS |
|  | 2503 / 2003 | 38.2 | 40.6 | 0.433 | BLOQ | 0.466 | BLOQ | BLOQ | 0.0940 | BLOQ | 0.563 |
|  | **Mean** | **66.3** | **61.7** | **0.433** | **0.248** | **0.653** | **0.247** | **BLOQ** | **0.124** | **BLOQ** | **0.544** |
|  | **SD** | **24.3** | **26.3** | **-** | **-** | **0.264** | **-** | **-** | **0.0424** | **-** | **0.0269** |
| Group 3 | 3501 / 3001 | 153 | 187 | 0.773 | 0.836 | BLOQ | 0.193 | BLOQ | BLOQ | 0.510 | 1.14 |
| 10 mg/kg | 3502 / 3002 | 110 | 146 | 0.410 | BLOQ | 0.500 | BLOQ | 0.483 | 0.117 | 0.324 | 0.183 |
|  | 3503 / 3003 | 128 | 193 | BLOQ | 2.46 | BLOQ | 1.16 | 0.290 | BLOQ | 0.121 | 0.178 |
|  | **Mean** | **130** | **176** | **0.591** | **1.65** | **0.500** | **0.678** | **0.387** | **0.117** | **0.318** | **0.499** |
|  | **SD** | **21.6** | **25.5** | **0.257** | **1.15** | **-** | **0.686** | **0.136** | **-** | **0.195** | **0.551** |
| Group 4 | 4501 / 4001 | 559 | 510 | 3.95 | 1.34 | 0.181 | 1.94 | 1.29 | BLOQ | 0.282 | BLOQ |
| 30 mg/kg | 4502 / 4002 | 379 | 524 | 1.11 | 3.49 | 1.54 | 2.12 | BLOQ | 1.00 | BLOQ | 0.857 |
|  | 4503 / 4003 | 356 | 265 | 1.50 | 2.20 | BLOQ | 0.396 | 0.273 | 1.41 | BLOQ | 0.256 |
|  | **Mean** | **431** | **433** | **2.19** | **2.34** | **0.862** | **1.48** | **0.781** | **1.21** | **0.282** | **0.557** |
|  | **SD** | **111** | **146** | **1.54** | **1.08** | **0.962** | **0.946** | **0.718** | **0.287** | **-** | **0.425** |

Supp Table 4: Distribution of MRG-229 Antisense strand to organ tissues in NHPs

| **Test Article** | **Dose (mg/kg)** | **Animal** | **Tissue Concentrations (ug/g)** | | | | |
| --- | --- | --- | --- | --- | --- | --- | --- |
|  |  |  | **Kidney** | **Liver** | **Spleen** | **Heart** | **Lung** |
| MRG-229 | 5 | 2001 | 16.1 | 4.75 | 0.184 | 0.147 | BQL |
|  |  | 2501 | 12.8 | 7.49 | 0.365 | BQL | BQL |
|  |  | **Mean** | **14.5** | **6.12** | **0.274** | **0.147** | **BQL** |
|  | 15 | 3001 | 32.4 | 14.6 | 2.63 | 0.131 | BQL |
|  |  | 3501 | 9.49 | 12.1 | 0.356 | 0.130 | BQL |
|  |  | **Mean** | **20.9** | **13.4** | **1.49** | **0.130** | **BQL** |
|  | 45 | 4001 | 18.4 | 21.5 | 2.44 | 0.809 | 0.335 |
|  |  | 4501 | 21.5 | 41.8 | 1.90 | 0.431 | 0.112 |
|  |  | **Mean** | **20.0** | **31.6** | **2.17** | **0.620** | **0.224** |

**Supp Table 5: Plasma Concentrations of MRG-229 Antisense Strand in Rats**

| **Group and Dose** | **PK Period** | **Animal** | **Sex** | **Concentration (µg/mL)** | | | | | | | |
| --- | --- | --- | --- | --- | --- | --- | --- | --- | --- | --- | --- |
|  |  |  |  | **Time (hr)** | | | | | | | |
|  |  |  |  | **0** | **0.083** | **0.5** | **1** | **2** | **4** | **8** | **24** |
| Group 3 | First Dose | 3504 | F | BLOQ | 15.2 | 3.01 | 1.77 | 1.65 | 0.875 | 0.116 | BLOQ |
| 10 mg/kg |  | 3505 | F | 0.00590 | 37.3 | 2.97 | 0.606 | 0.0745 | 0.0150 | 0.00650 | BLOQ |
|  |  | 3506 | F | BLOQ | 24.4 | 5.86 | 1.67 | 0.0581 | 0.00990 | 0.0189 | BLOQ |
|  |  |  | **Mean** | **0.00590** | **25.6** | **3.95** | **1.35** | **0.593** | **0.300** | **0.0470** | **BLOQ** |
|  |  |  | **SD** | **-** | **11.1** | **1.65** | **0.645** | **0.913** | **0.498** | **0.0597** | **-** |
| Group 3 | Last Dose | 3504 | F | BLOQ | 45.6 | 2.50 | 0.534 | 0.0306 | 0.00510 | BLOQ | 0.00910 |
| 10 mg/kg |  | 3505 | F | BLOQ | 47.0 | 2.73 | 0.806 | 0.0397 | 0.0196 | BLOQ | BLOQ |
|  |  | 3506 | F | BLOQ | 44.7 | 3.32 | 0.679 | 0.0274 | 0.0150 | BLOQ | BLOQ |
|  |  |  | **Mean** | **BLOQ** | **45.8** | **2.85** | **0.673** | **0.0326** | **0.0132** | **BLOQ** | **0.00910** |
|  |  |  | **SD** | **-** | **1.18** | **0.424** | **0.136** | **0.00638** | **0.00741** | **-** | **-** |
| Group 2 | Last Dose | 2001 | M | BLOQ | 15.9 |  |  |  |  |  |  |
| 3 mg/kg |  | 2002 | M | BLOQ | 17.2 |  |  |  |  |  |  |
|  |  | 2003 | M | 0.00580 | 15.0 |  |  |  |  |  |  |
|  |  | 2501 | F | BLOQ | 14.9 |  |  |  |  |  |  |
|  |  | 2502 | F | BLOQ | 13.5 |  |  |  |  |  |  |
|  |  | 2503 | F | BLOQ | 14.4 |  |  |  |  |  |  |
|  |  |  | **Mean** | **0.00580** | **15.2** |  |  |  |  |  |  |
|  |  |  | **SD** | **-** | **1.27** |  |  |  |  |  |  |
| Group 4 | Last Dose | 4001 | M | BLOQ | 99.9 |  |  |  |  |  |  |
| 30 mg/kg |  | 4002 | M | BLOQ | 121 |  |  |  |  |  |  |
|  |  | 4003 | M | BLOQ | 119 |  |  |  |  |  |  |
|  |  | 4501 | F | BLOQ | 129 |  |  |  |  |  |  |
|  |  | 4502 | F | BLOQ | 118 |  |  |  |  |  |  |
|  |  | 4503 | F | BLOQ | 116 |  |  |  |  |  |  |
|  |  |  | **Mean** | **BLOQ** | **117** |  |  |  |  |  |  |
|  |  |  | **SD** | **-** | **9.48** |  |  |  |  |  |  |

**Supp Table 6: Pharmacokinetic Parameters of TK Animals in Rats**

| **Group and Dose** | **Animal** | **First Dose** | | | | **Last Dose** | | | |
| --- | --- | --- | --- | --- | --- | --- | --- | --- | --- |
|  |  | **C_max_** | **AUC_last_** | **AUC_Back Ext_** | **t_1/2_** | **C_max_** | **AUC_last_** | **AUC_Back Ext_** | **t_1/2_** |
|  |  | **(µg/mL)** | **(µg*hr/mL)** | **(%)** | **(hr)** | **(µg/mL)** | **(µg*hr/mL)** | **(%)** | **(hr)** |
| Group 3 | 3510 | 15.2 | 11.4 | 12.8 | 1.53 | 45.6 | 12.3 | 40.4 | Und |
| 10 mg/kg | 3511 | 37.3 | 10.8 | 37.4 | 1.83 | 47.0 | 12.9 | 41.0 | Und |
|  | 3512 | 24.4 | 10.0 | 23.2 | Und | 44.7 | 12.6 | 38.7 | Und |
|  | **Mean** | **25.6** | **10.8** | **24.4** | **1.68** | **45.8** | **12.6** | **40.0** | **Und** |
|  | **SD** | **11.1** | **0.706** | **12.4** | **0.211** | **1.18** | **0.275** | **1.16** | **-** |

**Supp Table 7: Dose Proportionality of Last Dose Cmax Values in Rats**

| **Dose Levels Compared** | **Fold Change in Dose** | **Fold Change in C_max_** |
| --- | --- | --- |
| 3 mg/kg to 10 mg/kg | 3.3 | 3.0 |
| 10 mg/kg to 30 mg/kg | 3.0 | 2.6 |
| 3 mg/kg to 30 mg/kg | 10 | 7.7 |

**Supp Table 8: Male vs Female Cmax Values in Terminal Animal Groups**

| **Group and Dose** | **Animal #'s F/M** | **C_max_ (µg/mL)** | |
| --- | --- | --- | --- |
|  |  | **Female** | **Male** |
| Group 2 | 2501 / 2001 | 14.9 | 15.9 |
| 3 mg/kg | 2502 / 2002 | 13.5 | 17.2 |
|  | 2503 / 2003 | 14.4 | 15.0 |
|  | **Mean** | **14.3** | **16.0** |
|  | **SD** | **0.696** | **1.11** |
| Group 4 | 4501 / 4001 | 129 | 99.9 |
| 30 mg/kg | 4502 / 4002 | 118 | 121 |
|  | 4503 / 4003 | 116 | 119 |
|  | **Mean** | **121** | **113** |
|  | **SD** | **6.60** | **11.7** |

**Supp Table 9: Plasma Concentrations of MRG-229 Antisense Strand in NHPs**

| **Test Article** | **Dose (mg/kg)** | **Day** | **Animal** | **Concentration (µg/mL)** | | | | | | |
| --- | --- | --- | --- | --- | --- | --- | --- | --- | --- | --- |
|  |  |  |  | **Time (hr)** | | | | | | |
|  |  |  |  | **0.083** | **0.25** | **0.5** | **1** | **3** | **8** | **24** |
| MRG-229 | 5 | Day 1 | 2001 | 46.1 | 17.5 | 5.70 | 0.422 | 0.0161 | 0.00640 | 0.00590 |
|  |  |  | 2501 | 29.2 | 8.76 | 1.27 | 0.209 | 0.00970 | 0.00570 | BQL |
|  |  |  | **Mean** | **37.7** | **13.1** | **3.49** | **0.315** | **0.0129** | **0.00605** | **0.00590** |
|  |  | Day 15 | 2001 | 34.2 | 19.2 | 3.82 | 0.294 | 0.0308 | 0.0378 | 0.00580 |
|  |  |  | 2501 | 38.1 | 9.21 | 1.93 | 0.217 | 0.0114 | BQL | 0.00730 |
|  |  |  | **Mean** | **36.1** | **14.2** | **2.88** | **0.256** | **0.0211** | **0.0378** | **0.00655** |
|  | 15 | Day 1 | 3001 | 87.5 | 39.9 | 13.3 | 1.92 | 0.0781 | 0.0121 | BQL |
|  |  |  | 3501 | 105 | 27.6 | 7.74 | 0.937 | 0.0749 | 0.0143 | BQL |
|  |  |  | **Mean** | **96.1** | **33.8** | **10.5** | **1.43** | **0.0765** | **0.0132** | **-** |
|  |  | Day 15 | 3001 | 89.6 | 37.1 | 11.9 | 1.28 | 0.0689 | 0.0141 | 0.0112 |
|  |  |  | 3501 | 104 | 33.1 | 7.63 | 1.06 | 0.0482 | 0.0113 | BQL |
|  |  |  | **Mean** | **96.6** | **35.1** | **9.79** | **1.17** | **0.0586** | **0.0127** | **0.0112** |
|  | 45 | Day 1 | 4001 | 325 | 103 | 40.1 | 9.44 | 0.721 | 0.0865 | 0.0121 |
|  |  |  | 4501 | 298 | 105 | 32.3 | 5.50 | 0.277 | 0.0471 | BQL |
|  |  |  | **Mean** | **312** | **104** | **36.2** | **7.47** | **0.499** | **0.0668** | **0.0121** |
|  |  | Day 15 | 4001 | 350 | 134 | 36.1 | 8.88 | 0.540 | 0.0617 | 0.0124 |
|  |  |  | 4501 | 311 | 123 | 26.4 | 4.42 | 0.189 | 0.0198 | 0.0110 |
|  |  |  | **Mean** | **330** | **129** | **31.3** | **6.65** | **0.364** | **0.0408** | **0.0117** |

**Supp Table 10: Mean and Individual Pharmacokinetic Parameters in NHPs**

| **Test Article** | **Dose  (mg/kg)** | **Animal** | **Day 1** | | | | | | | **Day 15** | | | | | | |
| --- | --- | --- | --- | --- | --- | --- | --- | --- | --- | --- | --- | --- | --- | --- | --- | --- |
|  |  |  | **T_max_** | **C_max_** | **C_max_/D** | **AUC_last_** | **AUC_last_/D** | **R^2^** | **t_1/2_** | **T_max_** | **C_max_** | **C_max_/D** | **AUC_last_** | **AUC_last_/D** | **R^2^** | **t_1/2_** |
|  |  |  | **(hr)** | **(µg/mL)** | **(kg*µg/mL/mg)** | **(µg*hr/mL)** | **(hr*kg*µg/mL/mg)** |  | **(hr)** | **(hr)** | **(µg/mL)** | **(kg*µg/mL/mg)** | **(µg*hr/mL)** | **(hr*kg*µg/mL/mg)** |  | **(hr)** |
| MRG-229 | 5 | 2001 | 0.083 | 46.1 | 9.23 | 13.9 | 2.78 | 0.54 | UND | 0.083 | 34.2 | 6.83 | 11.4 | 2.27 | 0.90 | 7.82 |
|  |  | 2501 | 0.083 | 29.2 | 5.84 | 7.58 | 1.52 | 0.65 | UND | 0.083 | 38.1 | 7.62 | 9.88 | 1.98 | 0.44 | UND |
|  |  | **Mean** | **0.083** | **37.7** | **7.53** | **10.7** | **2.15** | **0.60** | **UND** | **0.083** | **36.1** | **7.23** | **10.6** | **2.12** | **0.67** | **7.82** |
|  | 15 | 3001 | 0.083 | 87.5 | 5.83 | 29.3 | 1.95 | 0.85 | 1.06 | 0.083 | 89.6 | 5.98 | 28.4 | 1.90 | 0.58 | UND |
|  |  | 3501 | 0.083 | 105 | 6.98 | 28.4 | 1.89 | 0.87 | 1.27 | 0.083 | 104 | 6.91 | 28.7 | 1.91 | 0.81 | UND |
|  |  | **Mean** | **0.083** | **96.1** | **6.41** | **28.8** | **1.92** | **0.86** | **1.16** | **0.083** | **96.6** | **6.44** | **28.6** | **1.90** | **0.70** | **-** |
|  | 45 | 4001 | 0.083 | 325 | 7.23 | 105 | 2.33 | 0.90 | 3.91 | 0.083 | 350 | 7.78 | 111 | 2.46 | 0.86 | 4.32 |
|  |  | 4501 | 0.083 | 298 | 6.61 | 90.3 | 2.01 | 0.86 | 1.12 | 0.083 | 311 | 6.90 | 91.7 | 2.04 | 0.66 | UND |
|  |  | **Mean** | **0.083** | **312** | **6.92** | **97.6** | **2.17** | **0.88** | **2.52** | **0.083** | **330** | **7.34** | **101** | **2.25** | **0.76** | **4.32** |

**Supp Table 11: C_max_ and AUC_last_ Dose Proportionality in NHPs**

| **Test Article** | **Doses Compared** | **Change in Dose** | **Change in C_max_** | | **Change in AUC_last_** | |
| --- | --- | --- | --- | --- | --- | --- |
|  |  |  | **Day 1** | **Day 15** | **Day 1** | **Day 15** |
| MRG-229 | 5 to 15 | 3x | 2.6 | 2.7 | 2.7 | 2.7 |
|  | 15 to 45 | 3x | 3.2 | 3.4 | 3.4 | 3.5 |
|  | 5 to 45 | 9x | 8.3 | 9.1 | 9.1 | 9.5 |
